# SMAD6 Integrates Endothelial Cell Homeostatic Flow Responses Downstream of Notch

**DOI:** 10.1101/2020.07.02.184820

**Authors:** Dana L Ruter, Ziqing Liu, Kimlynn M Ngo, X Shaka, Allison Marvin, Danielle B Buglak, Elise J Kidder, Victoria L Bautch

## Abstract

Laminar shear stress regulates blood vessel morphogenesis and subsequent quiescence, leading to vascular homeostasis. Although important for vessel function, how vascular homeostasis is set up and maintained is poorly understood. SMAD6, a scaffold for several signaling pathways, is expressed in developing arteries and its expression is flow-regulated. We found that SMAD6 is essential for endothelial cell flow-mediated responses downstream of the mechanosensor Notch1. Endothelial cells with reduced SMAD6 levels failed to align under homeostatic laminar shear flow, while forced SMAD6 expression rescued misalignment induced by reduced Notch1 signaling. SMAD6-dependent homeostatic laminar flow responses required the Notch ligand Dll4 and Notch transcriptional activity. Mechanistically, neither the N-terminal nor the C-terminal domain of SMAD6 alone rescued flow alignment upon loss of Notch signaling. Endothelial cells with reduced Smad6 levels had compromised barrier function, and RNA profiling revealed upregulation of proliferation-associated genes and down regulation of junction-associated genes. Among junction-related genes affected by SMAD6 levels, the proto-cadherin PCDH12 was upregulated by homeostatic flow and required for proper flow-mediated endothelial cell alignment. Thus, SMAD6 is a critical integrator of flow-mediated signaling inputs downstream of Notch1, as vessels transition from an angiogenic to a homeostatic phenotype.

## INTRODUCTION

Blood vessel formation intersects with tissue metabolic needs and architecture in multiple ways, including remodeling of the initial vascular plexus via shear stress provided by blood flow (Augustin and Koh, 2017; Carmeliet and Jain, 2011; Chappell et al., 2012). Endothelial cell responses orchestrate this remodeling and eventual transition to vascular homeostasis, as elevated heart pumping and blood volume increase the amount of fluid shear stress (Baeyens et al., 2016; Davies et al., 2003; Gimbrone et al., 2000; Hoffman et al., 2011). In contrast to the sprouting and blood vessel remodeling that characterize the “activated” endothelium of developmental stages, endothelial cells become quiescent under laminar flow at steady state. This is an important prerequisite for endothelial cell functions involved in homeostasis, including formation of a vascular barrier that regulates oxygen and nutrient exchange (Claesson-Welsh, 2015; Küppers et al., 2014; Lampugnani et al., 2018; Seebach et al., 2007). As homeostasis is induced, endothelial cells reduce proliferation and remodel their cytoskeleton and junctions to align the long axis parallel to flow, presumably to reduce flow-induced cellular stress (Dieterich et al., 2000; McCue et al., 2006; Noria et al., 1999; Schnittler et al., 1997; Shikata et al., 2005). The importance of this response is highlighted by the association of arterial laminar flow with protection from atherosclerosis, and the correlation between disturbed flow and atheroprone regions of vessels (Conway and Schwartz, 2013; Davies et al., 2010). Although several proteins and protein complexes have been identified as direct endothelial cell mechanosensors (Hahn and Schwartz, 2009), and even more signaling pathways have flow-responsive components, how these sensors and pathways integrate flow-mediated inputs remains poorly understood.

Notch signaling is implicated in flow-mediated changes in the vasculature, and recently Notch1 was identified as an endothelial cell mechanosensor (Mack et al., 2017; Polacheck et al., 2017). Endothelial cell deletion of Notch1 leads to flow misalignment, increased permeability, and increased atherosclerotic plaque formation. More generally, Notch signaling is upregulated in arteries that are exposed to higher shear stress than that found in veins of equivalent diameter, and Notch is thought to be responsible for maintaining arterial differentiation (Fang et al., 2017; Lawson et al., 2001; Pitulescu et al., 2017).

BMP signaling regulates several aspects of blood vessel formation and function in complex and context-dependent ways (Cai et al., 2012; Chappell et al., 2011; David et al., 2009; García de Vinuesa et al., 2016), with pro-angiogenic signaling downstream of ligands such as BMP2 and BMP6 countered by anti-angiogenic signaling downstream of BMP9 and/or BMP10. Human mutations in the Type 1 receptor Alk1/ACVRL1, the co-receptor Endoglin, or a common signaling component SMAD4 lead to Hereditary Hemmorrhagic Telangiectasia (HHT) that is characterized by arterio-venous malformations (AVMs) and hemorrhage (Gallione et al., 2010; Johnson et al., 1996; McAllister et al., 1994). Animal studies indicate that AVM formation requires genetic loss of BMP signaling, blood flow, and a third pro-angiogenic or pro-inflammatory stimulus (Garrido-Martin Eva et al., 2014; Roman and Hinck, 2017; Tual-Chalot et al., 2015; Urness et al., 2000), suggesting an intimate relationship between BMP signaling and flow responses in blood vessels.

There are several intersection points between Notch and BMP signaling in endothelial cells. For example, transcriptional responses are altered in complex and interdependent ways when both Notch and BMP signaling are activated, leading to cooperative effects on signaling and endothelial cell quiescence (Itoh et al., 2004; Larrivee et al., 2012; Rostama et al., 2015). We previously showed that pro-angiogenic endothelial cell BMP signaling *in vitro* and *in vivo* is regulated in part by Notch activation of an intracellular negative regulator of BMP signaling, SMAD6 (Mouillesseaux et al., 2016). SMAD6 expression is upregulated by laminar blood flow and elevated in arteries (Mouillesseaux et al., 2016; Topper et al., 1997; Wylie et al., 2018), suggesting that SMAD6 may function in vascular homeostasis. Here, we examined the role of SMAD6 in flow-mediated endothelial cell alignment and vascular homeostasis, which is the state of most healthy adult endothelium. We found that SMAD6 is required for flow-mediated homeostatic alignment of endothelial cells, and that SMAD6 expression rescues endothelial cell misalignment induced by loss of Notch signaling. RNA profiling revealed that endothelial cell-cell junction genes were down-regulated with reduced SMAD6 levels, and a proto-cadherin regulated downstream of SMAD6 affected endothelial cell flow alignment responses. Thus, SMAD6 is a critical integrator of homeostatic endothelial cell flow-mediated responses downstream of Notch signaling.

## RESULTS

### SMAD6 is Required for Homeostatic Laminar Flow-Induced Endothelial Cell Alignment and Polarization

Although endothelial cell responses to disturbed flow and changes with laminar flow have been investigated, the endothelial cell pathways and mechanisms that maintain vascular homeostasis once it is achieved are less well-understood. To begin to understand how endothelial cell flow-mediated homeostasis is maintained, we examined how the BMP regulator, SMAD6, regulates endothelial cell responses to extended periods of laminar shear stress. *Smad6*^*-/-*^ mutant embryos and pups with a knock-in *lacZ* reporter in the *Smad6* locus strongly express *lacZ* in endothelial cells of embryonic and early post-natal arteries, but not veins of similar diameter and stage, after the onset of blood flow (Galvin et al., 2000; Wylie et al., 2018). Since shear stress induced by laminar flow is higher in arteries, this suggests that endothelial SMAD6 expression is induced by homeostatic laminar flow. Thus, we chose conditions of 15 dynes/cm^2^ (d/cm^2^) for 72 hr for analysis of SMAD6 function in homeostatic arterial flow responses, and all output labeled “flow” is assessed from endothelial cells exposed to these conditions. Under these conditions, both human umbilical vein endothelial cells (HUVEC) and human arterial endothelial cells (HAEC) showed a 2-3 fold increase in Smad6 RNA (**Supp. Fig 1A-B**), consistent with a previous report (Topper et al., 1997).

**Figure 1.**
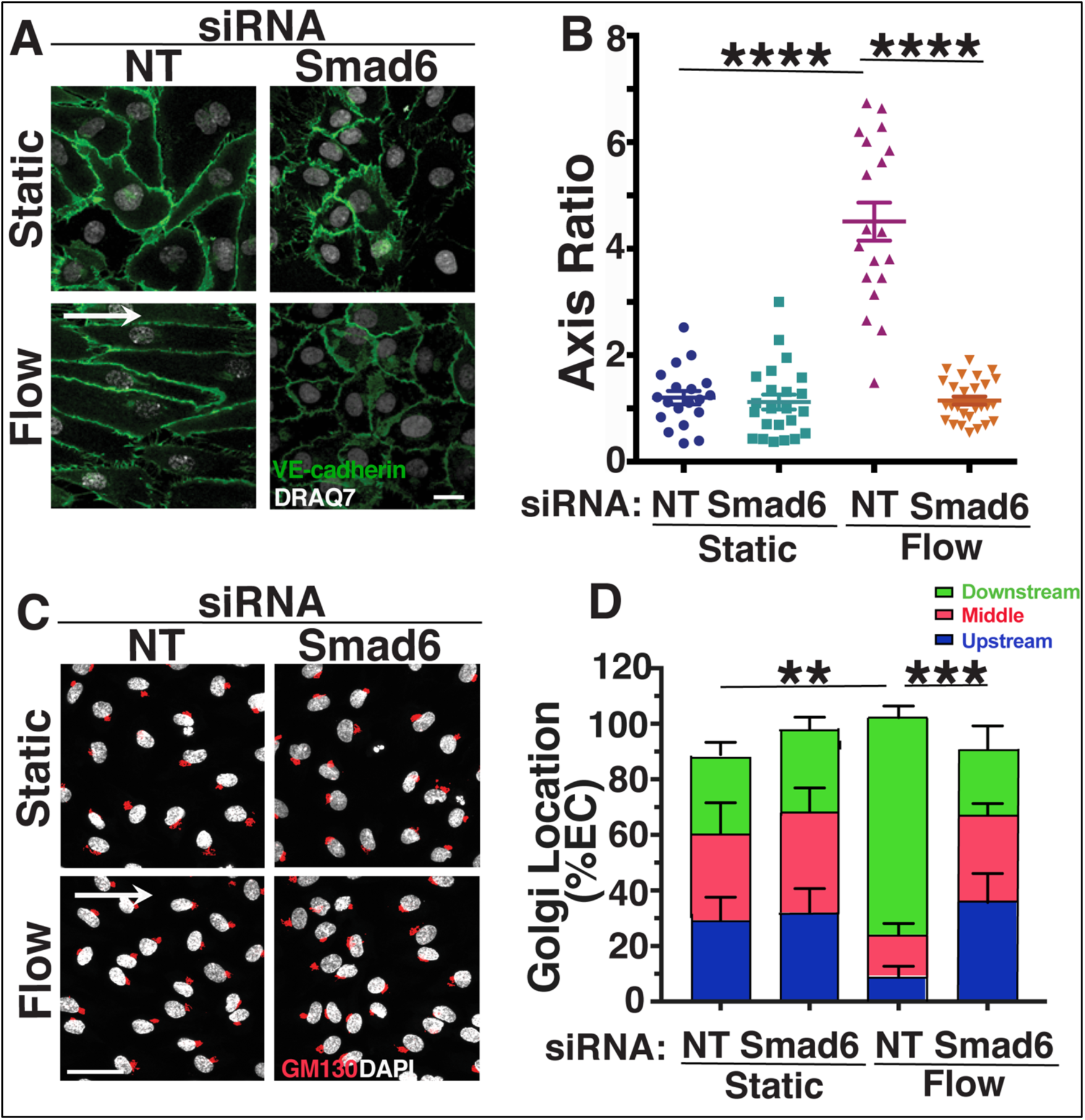
SMAD6 is Required for Homeostatic Endothelial Cell Flow-Mediated Alignment and Polarization. **A)** Representative panels of HUVEC stained with VE-cadherin (green, junctions) and DRAQ7 (white, nucleus) under static (no flow) or flow (laminar, 72h, 15 d/cm^2^ for all panels) conditions with indicated treatments. White arrow, flow vector. Scale bar, 20μM. **B)** Quantification of cell axis ratio in indicated conditions. Statistical analysis, One-way ANOVA; ****, p≤0.0001. **C)** Representative panels of HUVEC stained with GM130 (red, Golgi) and DAPI (white, nucleus) under indicated conditions with indicated treatments. White arrow, flow vector. Scale bar, 50μM. **D**) Quantification of Golgi localization relative to nucleus in indicated conditions. ≥30 cells/condition were analyzed. Statistical analysis, One-way ANOVA; **, p≤0.01; ***, p≤0.001. NT, non-targeting (control) siRNA.

We next asked whether SMAD6 is functionally important in endothelial cell responses to homeostatic laminar flow, by assessing HUVEC and HAEC exposed to flow and reduced levels of Smad6 RNA via knockdown (KD). We found that both venous and arterial endothelial cells with reduced Smad6 RNA levels failed to align with homeostatic laminar flow, as indicated by cell axis ratio and nuclear displacement angle (**Fig. 1A-B, Supp.Fig1C-F**). Endothelial cell misalignment in response to homeostatic laminar flow occurred with multiple siRNAs targeting Smad6 (Smad6-1 and Smad6-2), including a Smad6 siRNA pool (Smad6-3) (**Supp.Fig. 1G-H**). Both venous and arterial endothelial cells depleted for Smad6 also had mis-positioned Golgi and centrosomes under flow compared to controls (**Fig. 1C-D, Supp. Fig I-K, not shown**), indicating that SMAD6 is also important for endothelial cell polarization in response to homeostatic laminar flow. These results show that endothelial cell SMAD6 expression is flow-regulated and induced by homeostatic laminar flow, and that SMAD6 is functionally required for flow-mediated endothelial cell alignment and polarization.

### SMAD6 is Required Downstream of Notch for Flow-Mediated Alignment of Endothelial Cells

SMAD6 is both a negative regulator of BMP signaling and a transcriptional target of the pathway (Afrakhte et al., 1998; Imamura et al., 1997; Ishida et al., 2000; Larrivee et al., 2012). Consistent with the importance of canonical BMP signaling for endothelial cell flow alignment (Poduri et al., 2017), endothelial cells were misaligned under flow when incubated with the BMP inhibitor Crossveinless-2 (CV2) **(Supp. Fig 2A-B)**. However, since BMP receptors are not identified as direct mechanotransducers of flow-mediated signals, we searched for another pathway more directly linked to mechanotransduction that regulates SMAD6. In the absence of flow, Notch signaling upregulates SMAD6 expression, and Notch-induced SMAD6 regulates endothelial cell responsiveness to pro-angiogenic BMP ligands (Mouillesseaux et al., 2016). It was recently shown that Notch is a direct mechanotransducer of flow-mediated signaling (Mack et al., 2017; Polacheck et al., 2017), leading us to hypothesize that SMAD6 functions downstream of Notch in endothelial cell responses to homeostatic laminar flow. Reduced Notch signaling, via siRNA depletion of Notch1 or by treatment with the γ-secretase inhibitor DAPT which blocks Notch signaling, prevented endothelial cell alignment under conditions of homeostatic laminar flow (**Fig. 2A-B, Supp. Fig 2C-D**). The misalignment induced by reduced Notch was accompanied by reduced expression of Smad6 RNA relative to controls **(Fig. 2C, F**), indicating that SMAD6 expression levels are regulated downstream of Notch in endothelial cells and are important for homeostatic flow alignment. We next asked whether ectopic SMAD6 expression rescued the flow-mediated misalignment of endothelial cells downstream of reduced Notch. Upon incubation with DAPT, control endothelial cells transfected with empty vector remained misaligned when exposed to homeostatic laminar flow and did not differ from nearby untransfected cells; in contrast, similarly treated endothelial cells expressing SMAD6 aligned in response to homeostatic laminar flow while nearby untransfected cells remained misaligned **(Fig 2D-E)**. Endothelial cells expressing SMAD6 after Notch1 KD also aligned to homeostatic laminar flow while nearby cells remained misaligned **(Fig 2G-H)**. Thus SMAD6 expression rescued Notch loss-induced endothelial cell misalignment, indicating that SMAD6 is an effector of Notch-mediated homeostatic flow alignment in endothelial cells.

**Figure 2.**
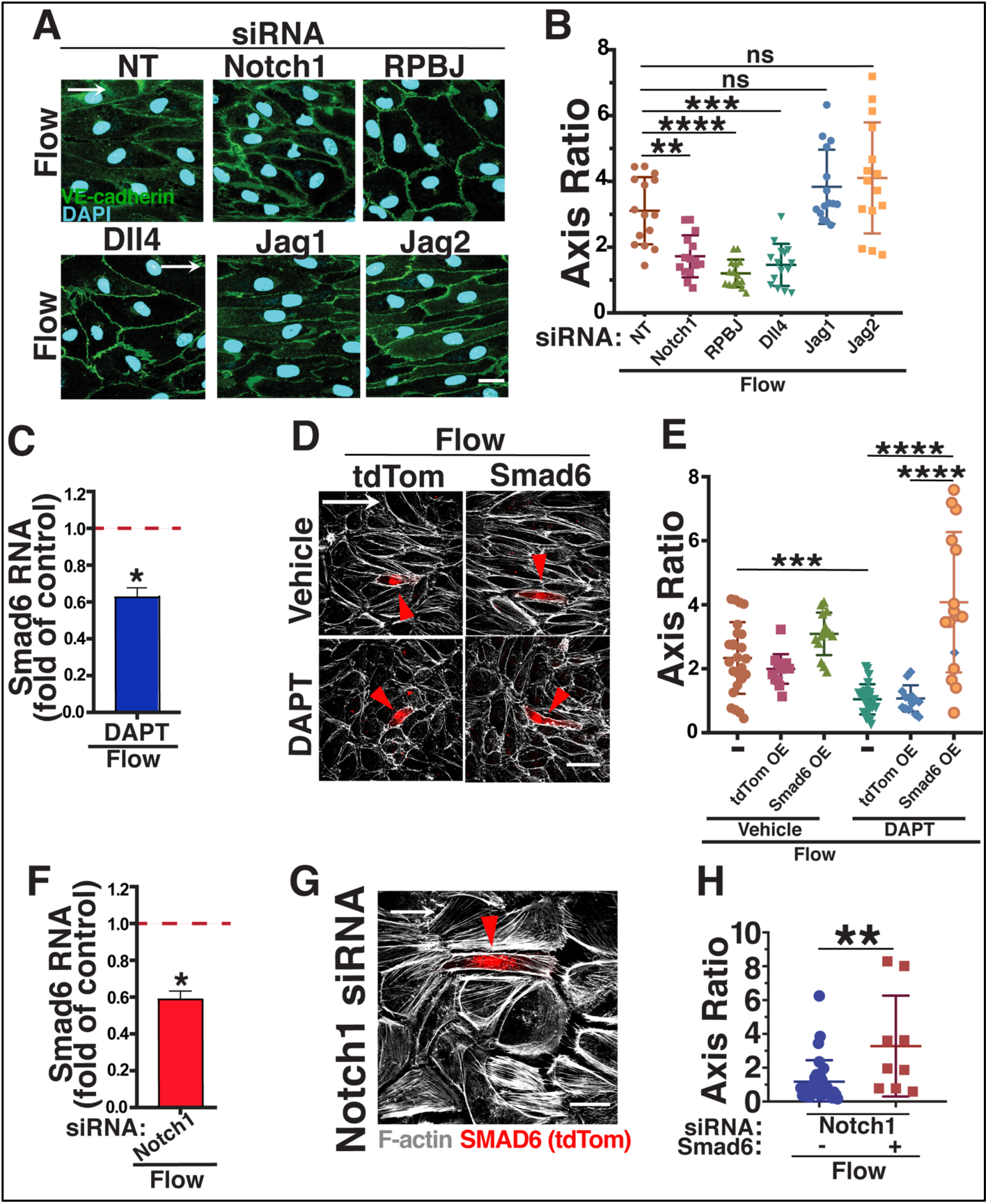
SMAD6 is Downstream of Notch Signaling in Homeostatic Endothelial Cell Flow-Mediated Alignment. **A**) Representative panels of HUVEC stained with VE-cadherin (green, junctions) and DAPI (blue, nucleus) under control (static) or flow conditions with indicated treatments. White arrow, flow vector. Scale bar, 20μM. **B)** Quantification of cell axis ratio. Statistical analysis, One-way ANOVA; **, p≤0.01; ***, p≤0.001; ****, p≤0.0001. **C)** qPCR RNA levels (normalized to vehicle control under flow) of SMAD6 in HUVEC treated with DAPT under flow. Statistical analysis, Student’s t-test; *, p≤0.05. **D**) Representative panels of HUVEC stained with Phalloidin (F-actin, white) with indicated treatments and expression constructs (Empty Vector (EV) or SMAD6) under flow conditions. White arrow, flow vector. Red arrowhead, positive HUVEC. Scale bar, 50μM. **E)** Quantification of cell axis ratio in indicated conditions. Statistical analysis, One-way ANOVA; ***, p≤0.001; ****, p≤0.0001. **F)** qPCR RNA levels (normalized to vehicle control under flow) in in HUVEC treated with Notch1 siRNA under flow. Statistical analysis, Student’s t-test; *, p≤0.05. **G)** Representative panels of HUVEC stained with Phalloidin (F-actin, white) with indicated treatments and expression construct (SMAD6) under flow conditions. White arrow, flow vector. Red arrowhead, positive EC. Scale bar, 20μM. **H)** Quantification of cell axis ratio in indicated conditions. Statistical analysis, One-way ANOVA; **, p≤0.01.

We next asked which Notch signaling components are required for homeostatic endothelial cell flow alignment. RPBJ is a transcriptional co-activator required for canonical downstream Notch signaling, and endothelial cells with reduced levels of RPBJ had reduced expression of Smad6 RNA **(Supp. Fig. 2E)**. Moreover, RPBJ knockdown led to misalignment of both arterial and venous endothelial cells in response to homeostatic laminar flow **(Fig. 2A-B; Supp. Fig 2F-I)**. Several Notch ligands are expressed in endothelial cells and implicated in Notch responses to flow. Reduced levels of Dll4, but not Jagged1 or Jagged2, resulted in endothelial cells misaligned in response to homeostatic laminar flow (**Fig. 2A-B; Supp.Fig 2F-G**), suggesting that Dll4-mediated activation of Notch1 signaling is important for homeostatic endothelial cell flow alignment via canonical Notch signaling.

The misalignment induced by RPBJ or Dll4 knockdown was rescued by expression of SMAD6 in HUVEC **(Fig. 3A-B)**, and SMAD6 expression also rescued alignment of arterial endothelial cells with homeostatic laminar flow after RPBJ KD **(Supp. Fig 2H-I)**, suggesting that Notch regulation of SMAD6 expression is an important component of endothelial cell responses to homeostatic flow. SMAD6 is comprised of two parts connected by a linker **(Supp Fig.2J)**; the N-terminal portion includes several arginine residues that are methylated to regulate SMAD6 activity (Xu et al., 2013), while the C-terminal portion contains the MH2 protein-interacting domain (Hanyu et al., 2001). Since both parts are required for the regulatory role of SMAD6 (Nakayama et al., 2001), we hypothesized that SMAD6 rescue of homeostatic endothelial cell alignment downstream of Notch required full-length SMAD6. Reduced Notch signaling, either via Notch1 or RPBJ depletion, led to endothelial cell misalignment in response to homeostatic flow that was not rescued by expression of constructs encoding only either the N-terminal or C-terminal portion of SMAD6 (**Fig. 3C-F)**. These data indicate that full-length SMAD6 is required to mediate the effects of Notch signaling on homeostatic endothelial cell flow alignment.

**Figure 3.**
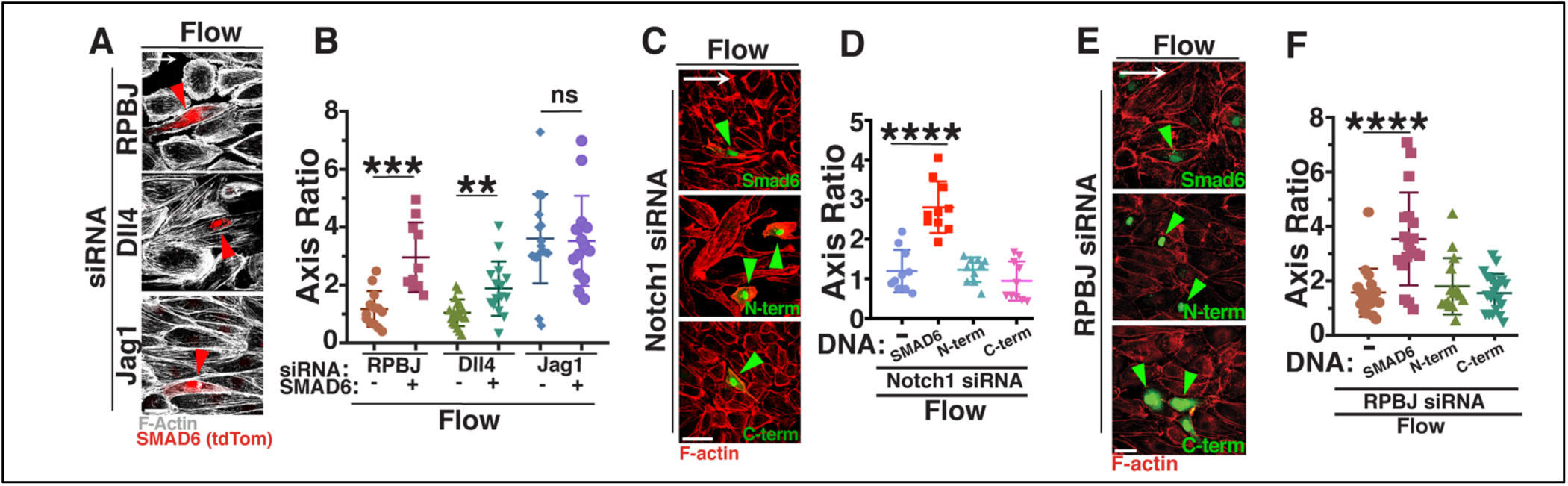
Full-Length SMAD6 Rescues Notch Loss-Induced Endothelial Cell Homeostatic Flow Mis-Alignment. **A)** Representative panels of HUVEC stained with phalloidin (F-actin, white) with indicated siRNA treatments and SMAD6 expression construct under flow conditions. White arrow, flow vector. Red arrowheads, positive EC. Scale bar, 50μM. **B)** Quantification of cell axis ratio in indicated conditions. Statistical analysis, One-way ANOVA; **, p≤0.01; ***, p≤0.001; ns, not significant. **C)** Representative panels of HUVEC stained with Phalloidin (F-actin, red) with Notch1 siRNA treatments and expression constructs (full-length SMAD6, N-terminal SMAD6, or C-terminal SMAD6; green) under flow conditions. White arrow, flow vector. Green arrowhead, positive EC. Scale bar, 50μM. **D)** Quantification of cell axis ratio in indicated conditions. Statistical analysis, One-way ANOVA; ****, p≤0.0001. **E)** Representative panels of HUVEC stained with Phalloidin (F-actin, red) with RPBJ siRNA treatment and expression constructs (full-length SMAD6, N-terminal SMAD6, or C-terminal SMAD6; green) under flow conditions. White arrow, flow vector. Green arrowhead, positive EC. Scale bar, 50μM. **F)** Quantification of cell axis ratio in indicated conditions. Statistical analysis, One-way ANOVA; ****, p≤0.0001.

### SMAD6 Regulates Endothelial Cell Proliferation

To better understand the effects of reduced SMAD6 function on flow-mediated endothelial cell responses, we examined the transciptome of endothelial cells under homeostatic laminar flow relative to non-flow conditions, and with depleted Smad6 levels. Pearson Correlation Analysis revealed good correlation between experimental replicates of each condition **(Supp. Fig 3A)**. Principle Component Analysis (PCA) distinguished the transcriptomes of control (non-targeting siRNA) and SMAD6 depleted endothelial cells, and transcriptomes also clustered by flow status **(Supp. Fig 3B)**. Overall comparisons (33,694 genes) showed that, when binned by flow status, only 1.2% of transcripts were significantly up- or down-regulated with reduced Smad6 levels compared to NT siRNA controls under static (non-flow) conditions, while 6.9% of transcripts changed with reduced Smad6 levels under homeostatic flow conditions **(Table 1)**. When binned by depletion condition, NT siRNA static vs. homeostatic laminar flow conditions led to 6.7% of transcripts significantly changing expression levels, while comparison after Smad6 KD showed that 8.9% of transcripts changed with flow compared to static conditions. These numbers suggest that the magnitude of flow-mediated changes is greatest in endothelial cells with reduced Smad6 levels, consistent with a role for SMAD6 in endothelial cell flow responses.

**Table 1.**
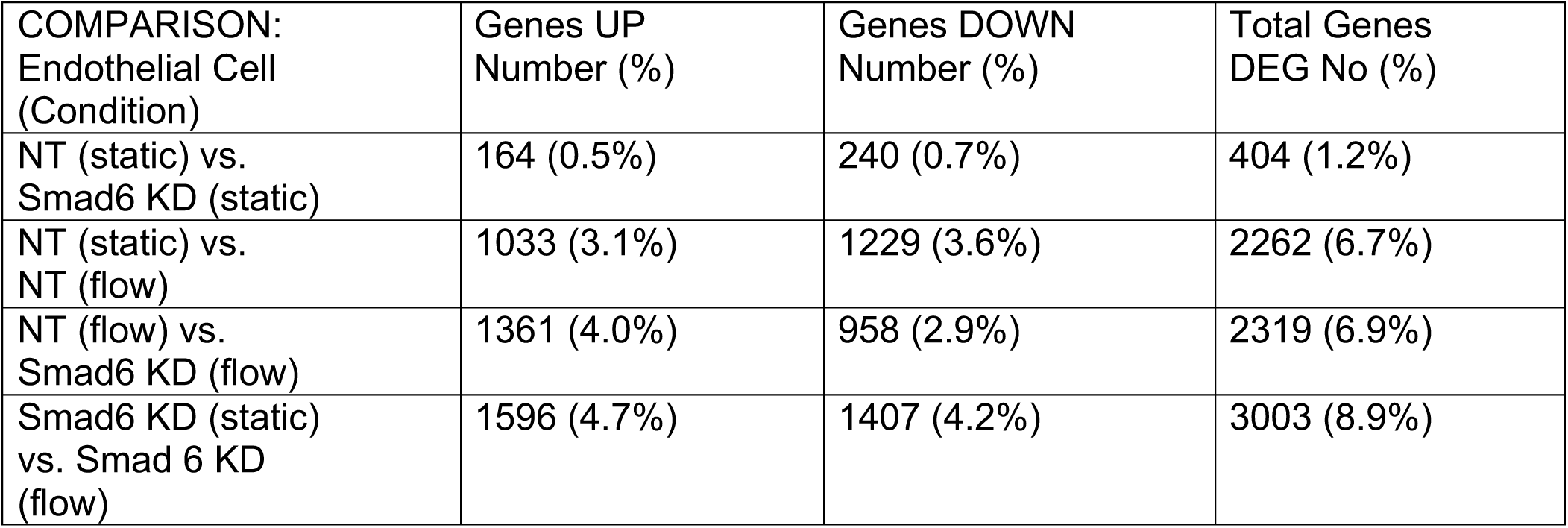
Changes in HUVEC Gene Expression (total 33,694).

| COMPARISON:<br>Endothelial Cell<br>(Condition) | Genes UP<br>Number (%) | Genes DOWN<br>Number (%) | Total Genes<br>DEG No (%) |
| --- | --- | --- | --- |
| NT (static) vs.<br>Smad6 KD (static) | 164 (0.5%) | 240 (0.7%) | 404 (1.2%) |
| NT (static) vs.<br>NT (flow) | 1033 (3.1%) | 1229 (3.6%) | 2262 (6.7%) |
| NT (flow) vs.<br>Smad6 KD (flow) | 1361 (4.0%) | 958 (2.9%) | 2319 (6.9%) |
| Smad6 KD (static)<br>vs. Smad 6 KD<br>(flow) | 1596 (4.7%) | 1407 (4.2%) | 3003 (8.9%) |

We next used Gene Ontogeny (GO) Analysis to examine gene categories that were significantly affected by Smad6 depletion, and we found that transcripts associated with the cell cycle and mitosis were enriched in endothelial cells with reduced Smad6 levels, independent of flow status **(Supp Fig. 3C)**. Thus we hypothesized that endothelial cell proliferation is negatively regulated by SMAD6. Consistent with this hypothesis, expression of the proliferation marker Ki67 was increased upon loss of Smad6 under both static and homeostatic flow conditions, and BrdU incorporation, which labels S-phase cells, was increased with reduced Smad6 in static conditions and trending upward under flow conditions **(Supp Fig. 3D-G**). However, endothelial cells with reduced Smad6 levels had reduced BrdU incorporation and Ki67+ reactivity under flow compared to static conditions, suggesting that the changes to endothelial cell proliferative capacity remain flow-responsive despite reduced SMAD6 levels.

### SMAD6 Regulates Endothelial Cell Barrier Function and Junctions

RNA profiling also indicated that genes associated with cell-cell junctions were down-regulated in endothelial cells with reduced Smad6 levels **(Fig. 4A)**, and we previously showed that SMAD6 regulates junction morphology in the absence of flow (Wylie et al., 2018). This led us to ask whether the barrier formed by endothelial cell-cell junctions and important for proper vascular function was compromised with loss of SMAD6. Endothelial cells with depleted Smad6 levels had reduced barrier function relative to controls, as measured by trans-endothelial electrical impedance **(Fig. 4B)**, and the abrogated resistance downstream of reduced Smad6 levels was also significant under flow conditions **(Fig. 4C)**.

**Figure 4.**
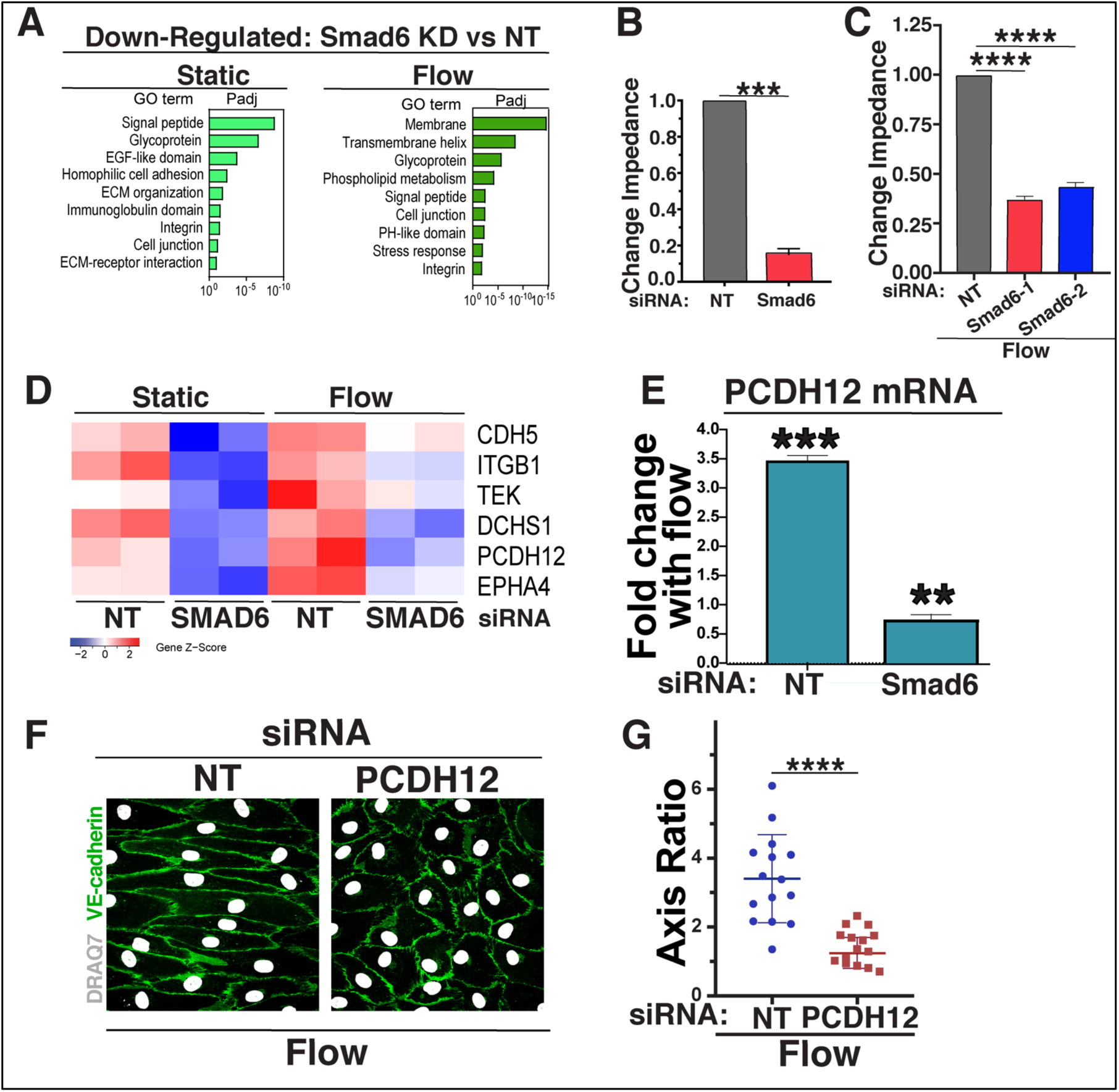
SMAD6 Regulates Endothelial Cell Barrier Function and Cell-Cell Junction Genes under Homeostatic Flow. **A**) Gene Ontology (GO) analysis performed on differentially expressed genes from bulk RNA-seq data sets using DAVID. GO terms significantly enriched (P adjusted <0.1) in down-regulated are shown. **B)** Change in impedence after 24 hr of EC with indicated siRNAs under static (control) conditions, normalized to control. Statistical analysis, One-way ANOVA; ***, p≤0.001. **C**) Change in impedence after 72 hr of EC with indicated siRNAs under flow conditions, normalized to control. Statistical analysis, One-way ANOVA; ****, p≤0.0001. **D)** Heat map showing subset of cell-cell-adhesion genes (see text for details) down-regulated with reduced Smad6 levels under static (control) and flow conditions. **E)** qPCR of relative levels of PCDH12 RNA under flow relative to static (control) with indicated siRNAs. **F**) Representative panels of HUVEC stained with VE-cadherin (green, junctions) and DAPI (white, nucleus) under control (static) or flow conditions with indicated treatments. White arrow, flow vector. Scale bar, 50μM. **G)** Quantification of cell axis ratio. Statistical analysis, student’s t-test; ****, p≤0.0001.

To further examine how SMAD6 manipulations affect endothelial cell-cell junctions, we focused on expression differences in cell-cell adhesion and junction genes between control and Smad6 KD endothelial cells under both static and flow conditions, Overlapping genes from these two genelists were identified, and a small subset with expression levels above 50 counts per million was examined further. Most junction genes were upregulated with homeostatic laminar flow in controls, and relative expression of a subset of these genes was reduced in endothelial cells depleted for Smad6, regardless of flow status. The net effect was that a group of cell junction genes had reduced expression in SMAD6-depleted endothelial cells relative to contols when both were exposed to homeostatic laminar flow **(Fig. 4D, compare Flow-NT to Flow-Smad6 KD)**.

Of these, one with significant expression changes that were both flow and SMAD6-dependent was chosen for further analysis. PCDH12 is a proto-cadherin expressed in arterial endothelial cells, and its deletion leads to changes in murine arterial blood pressure, while human mutations are associated with arterial calcification in the brain (Nicolas et al., 2017; Philibert et al., 2012; Rampon et al., 2005). First, we validated the expression changes in PCDH12 with qRT-PCR; we found that PCDH12 expression was significantly upregulated by homeostatic laminar flow, and this increase was blunted to 60% of static control levels in flowed endothelial cells with reduced Smad6 levels **(Fig. 4E)**. We next asked whether PCDH12 functions in flow alignment responses, by subjecting HUVEC treated with PCDH12 siRNA to homeostatic laminar flow, and found that PCDH12 depleted endothelial cells failed to align **(Fig. 4F-G)**. These results show that SMAD6 is required for full PCDH12 expression in response to flow, and that PCDH12 expression is necessary for proper endothelial cell alignment under homeostatic flow conditions. These findings suggest that SMAD6 regulation of PCDH12 contributes to endothelial cell homeostatic flow-mediated responses downstream of Notch signaling.

## DISCUSSION

This work reveals a functional requirement for SMAD6 in endothelial cell alignment and quiescence in response to homeostatic laminar flow, which is the shear stress experienced by arterial endothelial cells and is considered atheroprotective. SMAD6 is downstream of Notch1, and since Notch1 is a mechanotransducer, it is likely that shear stress signals, transduced by Notch1 signaling, mediate flow responses in part through SMAD6. Expression profiling reveals numerous SMAD6-dependent changes in gene expression, with predominant increases in cell cycle/proliferation genes and prominent down-regulation of cell-cell adhesion genes with reduced Smad6 levels, consistent with a role for SMAD6 in regulating endothelial cell quiescence in response to laminar flow. Thus SMAD6 is a critical integrator of Notch-dependent endothelial cell flow responses, and endothelial cell responses to flow that promote vascular homeostasis.

It is likely that Notch regulation of homeostatic flow responses is linked to its regulation of SMAD6 expression levels. Loss of Notch1 signaling significantly reduced Smad6 RNA levels under homeostatic flow conditions, and restored expression of SMAD6 was sufficient to rescue flow-mediated alignment downstream of loss of Notch signaling. Interestingly, the Notch requirement for endothelial cell alignment under homeostatic flow is dependent on the ligand Dll4 and requires Notch-mediated transcription, as both reduced levels of Dll4 and RPBJ led to endothelial cell misalignment that was rescued by restoring SMAD6 expression. It appears that initial endothelial flow-mediated responses do not require Notch transcriptional activity (Polacheck et al., 2017), although subsequent flow responses may depend on Notch transcriptional activity (Mack et al., 2017), suggesting that equilibration to laminar flow involves a switch from a non-transcriptional to a transcriptional program. Careful analysis of endothelial cell junctional changes to laminar flow indicate immediate effects on VE-cadherin clustering that are followed by later junction and cell shape changes (Seebach et al., 2007), consistent with the idea of a switch. A switch model is also consistent with SMAD6 expression regulation being critical for vascular homeostasis in response to flow, and suggests that endothelial cells may have evolved different mechanisms for an acute response to changes in mechanotransduction vs. equilibration to ongoing mechanotransduction inputs. We previously showed that sequences upstream of the SMAD6 gene contain binding sites for RBPJ (Mouillesseaux et al., 2016), so it is possible that these sequences are also involved in flow-mediated Notch regulation of SMAD6 expression.

Although SMAD6 functions as a negative regulator of BMP signaling, it also has BMP-independent functions in innate immunity and other cellular processes (Choi et al., 2006; Lee et al., 2015; Zhang et al., 2018), so it may affect endothelial cell flow responses in BMP-independent ways. Alternatively, canonical BMP signaling also affects flow responses in complex ways, so SMAD6 may regulate flow responses downstream of both Notch and BMP inputs, and it will be interesting to determine whether SMAD6 is a critical integrator of these complex pathways under flow. In any case, expression profiling showed that more genes change expression under homeostatic flow conditions when SMAD6 levels are reduced (6.9%) vs. non-flow conditions (1.2%), indicating that homeostatic flow amplifies SMAD6-dependent transcriptional differences in endothelial cells. Loss of SMAD6 prevents alignment of endothelial cells under flow, resulting in morphological differences that may account for some of the changes. However, comparison of endothelial cells with reduced SMAD6 levels and similar morphology in non-flow vs. flow conditions reveals an 8.9% difference in gene expression profiles, suggesting that many SMAD6-dependent transcription changes derive from reduced SMAD6 levels in response to flow, and are not secondary to changes in cell morphology.

The loss of SMAD6 led to a more “activated” endothelial cell phenotype, with cell cycle/proliferation pathways upregulated and cell-cell junction pathways down-regulated, and these changes were accompanied by significant loss of barrier function. These findings are consistent with SMAD6 regulating the atheroprotective endothelial cell quiescence phenotype that accompanies flow-mediated alignment under homeostatic flow and important for barrier function, and it is also consistent with the hemorrhage phenotype of *Smad6*^*-/-*^ mutant embryos (Wylie et al., 2018). Interestingly, cell cycle pathways were also upregulated in endothelial cells with reduced levels of Notch1 (Mack et al., 2017), consistent with SMAD6 being downstream of Notch activation. Although the profiling suggests that endothelial cell-cell adhesion changes with reduced SMAD6 are likely to be complex and involve multiple adhesion receptors, we found that the proto-cadherin PCDH12 is significantly upregulated by homeostatic laminar flow, and this upregulation was dramatically blunted by reduced Smad6 levels. Since independent reduction of PCDH12 levels abrogates endothelial cell flow-mediated alignment, it is likely that PCDH12 is one SMAD6 target normally upregulated downstream of SMAD6 under homeostatic flow that contributes to endothelial cell barrier function and quiescence. Thus we provide evidence that homeostatic flow-mediated mechanotransduction from Notch1 involves transcriptional activity that regulates expression of the effector SMAD6, and SMAD6 levels affect endothelial cell alignment, cell-cell junctions and barrier function. These findings provide new intersections and potential new therapeutic targets for diseases such as atherosclerosis that are linked to loss of vascular homeostasis.

## MATERIALS AND METHODS

### Cell Culture

HUVEC (Lonza, #C2519A) and HAEC (Lonza, #C2535) were maintained according to manufacturer’s recommendations and used at passage 2-4. For HUVEC and HAEC culture, EBM-2 (Endothelial Cell Growth Basal Medium-2, Lonza, #CC-3156) was supplemented with EGM-2 SingleQuots Supplements (Lonza, #CC-4176) (called EBM-2+). Main experiments were independently replicated with a different lot of HUVEC, and key experiments were replicated in HAEC at least two independent times. HUVEC and HAEC were certified mycoplasma-free by the UNC Tissue Culture Facility.

### Endothelial Cell Flow Experiments

HUVEC and HAEC were plated at 100% confluency in each lane of a µ-Slide VI^0.4^ (Ibidi, #80601) coated with fibronectin (Millipore Sigma, F2006, 5μg/mL) 4 hr prior to the experiment in EGM-2+ medium. After 2 hr for cell attachment and spread, slides were washed 3X in flow medium (EBM-2 with 10% FBS (Gibco, #26140-079) and 1X Antibiotic-Antimycotic (Gibco, #15240-062)), and incubated in flow medium for 2 hr prior to flow onset. Uniform laminar shear stress was generated by attaching slide chambers to a pump system (Ibidi, #10902) with both the slide and the pump apparatus kept at 37°C and 5% CO_2_. To reduce cell shearing, flow was applied for 30 min at 5 d/cm^2^, then 30 min at 10 d/cm^2^, followed by 72 hr at 15 d/cm^2^. For siRNA knockdown, siRNA incubation was for 30 hr prior to plating and flow initiation.

### Endothelial Cell Shear Stress Measurements

All data presented under “flow” conditions is at 15 d/cm^2^ laminar flow for 72 hr (Ibidi system) and considered to be homeostatic laminar flow.

#### Cell Axis Ratio

Cell shape and alignment were measured by staining for VE-cadherin or PECAM1 as described below. Each cell size measurements were recorded by measuring total cell length and width using ImageJ. Length divided by width was used for cell axis ratio. At least 10 cells/condition were measured, and each experiment was replicated at least 3 times.

#### Nuclear Displacement Angle

Nuclear displacement angle was measured using the angle measurement tool in Image J. Measured angles were degrees separating a line perpendicular to the flow vector and a line through the nucleus along its long axis **(see Supp. Fig. 1C)**. At least 10 cells/condition were measured, and each experiment was replicated at least 3 times.

#### Cell Polarization

We determined the Golgi or centrosome location relative to the nucleus. HUVEC and HAEC stained for the Golgi (GM130) or the centrosome (TUBGCP2) and nuclei (DAPI) were analyzed by dividing the nucleus into equal thirds and and binning the organelle relative to the flow vector: upstream, middle, or downstream. At least 5 fields/experiment were quantified per condition.

### siRNA Transfection

HUVEC (Lonza) were transfected with non-targeting siRNA (NT, Life Technologies, #4390847) or experimental siRNAs (single siRNAs or siRNA pools – see Key Resources) using the standard Lipofectamine 2000 (Invitrogen, 11668027) manufacturer’s protocol. Briefly, for each siRNA to be tested: 24 μl of 10uM siRNA was diluted in 476 μl of opti-MEM media and separately, 24 μl of Lipofectamine 2000 reagent was diluted in 476 μl of opti-MEM medium. Each mixture was incubated separately 5 min at RT, then mixed 15 min at RT. This mixture was added to a 10cm plate of ∼70% confluent HUVEC or HAEC in EGM-2+ medium without antibiotics (EBM-2 media, added BulletKit without gentamicin supplement) and incubated 24 hr at 37 °C, then incubated for 6 hr in fresh non-antibiotic EGM-2+ prior to plating in flow channels to start the experiment.

### Immunofluorescence

HUVEC and HAEC were fixed for 10 min in 4% PFA, washed 3X with PBS, then permeabilized for 10 min in 0.1% Triton X-100. Cells were blocked for 30 min at RT in 1% BSA (Millipore Sigma, #A-4503) in PBS, then incubated with primary antibodies (1:100—see Key Resources) in 1% BSA for 45 min at RT. After washing 5X with 1% BSA, samples were incubated with Alexa-fluor-conjugated anti-species secondary antibodies (1:250—see Key Resources) plus Alexa-fluor-conjugated phalloidin (1:50— Invitrogen, #A12379 or #A12381) and DAPI or DRAQ7 (1:300—Sigma #10236276001 or Abcam #ab109202, respectively) in 1% BSA for 30 min at RT. Samples were washed 5X in 1% BSA and mounted by washing 3X in 80% glycerol (Millipore Sigma, #G5516) in PBS. Immunofluorescent imaging was done either immediately after mounting or slides were wrapped in foil at 4° for up to 2 weeks.

BrdU incorporation was an adaptation (Yu et al., 2017). Briefly, HUVEC were incubated with 10 μM BrdU (Millipore Sigma, #B5002) in flow media for the last 90 min of flow and fixed in ice-cold 100% methanol. Samples were blocked for 30 min at RT in 1% BSA, then acid treated as follows: 10 min in cold (4°) 1N HCl on ice, rinse in 2N HCl and 10 min at RT, then incubated in 10mM citric acid solution (pH 7.4, in 0.2M Na^+2^HPO_4_) for 10 min at RT, rinsed with 1% BSA 3X and processed for antibody staining as detailed above with a sheep polyclonal antibody to BrdU (1:100—Abcam, #ab1893), then Alexa Fluor Donkey anti-sheep 594 secondary (1:250—Life Technologies #A-11016), along with DAPI and phalloidin as described above.

All fluorescent imaging was done using an Olympus FV3000 Laser Scanning Confocal Microscope and Flow View software. Olympus OIB file formats were imported into ImageJ using Bio-Formats Importer20 for analysis and quantification.

### Quantitative RT-PCR

Primers are listed in Key Resources at end of this section. cDNA was generated from 1 μg mRNA using iScript reverse transcription kit (Bio-Rad, #1708891) and diluted 1:3 in water. qRT–PCR was performed using iTaq Universal SYBR Green SuperMix (Bio-Rad, #1725121). SYBR Green real-time PCR was performed in triplicate on the Applied Biosystems QuantStudio 6 Flex Real-Time PCR System. For quantification, relative expression of each gene to GAPDH in each sample was calculated by 2^(CT of gene−CT of GAPDH). Statistical significance was determined by one-sample *T*-test compared to a reference value of onefold change.

### RNA Sequencing and Analysis

RNA was extracted using TRIzol (Invitrogen) from two biological replicates (independent experiments), and TruSeq Stranded mRNA Library Prep Kit (Illumina) was used to prepare cDNA and Illumina libraries for sequencing (HiSeq4000). 2-5 × 10^7^ 50-bp paired-end reads per sample were obtained and mapped to human genome GRCh38-1.2.0 downloaded from http://cf.10xgenomics.com/supp/cell-exp/ with TopHat/2.1.1 using default settings. Mapping rate was >92% for all samples, and gene expression was determined with Htseq-count/0.6.1 using the union mode (http://www-huber.embl.de/users/anders/HTSeq). PCA analysis was performed with the top 400 genes selected by largest weight (loading) contribution to PCs 1, 2 or 3 using the R package SINGuLAR. Differential expression analysis was performed with DESeq2 in R, and lists of differentially expressed genes were obtained (FDR < 0.05). Heat maps were generated using the heatmap.2 function in the ‘gplots’ package in R. Gene ontology analysis was performed using the DAVID functional annotation tool version 6.8 (https://david.ncifcrf.gov/). All gene ontology terms shown in this study have a corrected P value (the “Benjamini” value from DAVID) < 0.1. The RNA-seq data that support the findings of this study are available in the Gene Expression Omnibus (GEO) under the accession number GSE147036.

### Barrier Function Analysis

#### Real-Time Cell Analysis (RTCA) Experiments

Barrier properties were measured using a commercially available system (xCELLigence Real-Time Cell Analyzer (RTCA)); Acea Biosciences/Roche Applied Science, Basel, Switzerland). RTCA measures electrical impedance as a readout for the barrier status of cells grown on top of microelectrode coated surfaces. HUVEC were pre-treated with siRNAs or drug for 24 hr prior to plating an equal cell number onto the microelectrode surface of the E-plate (E-plate 16, Roche Applied Science). Impedance readings were taken automatically every 5 min for 24 hr.

#### Electric Cell-Substrate Impedance Sensing System (ECIS) Experiments

Endothelial barrier function analysis under static and flow conditions was performed using impedance-based cell monitoring using ECIS zeta theta (Applied Biophysics) in conjunction with the Ibidi pump system. HUVEC were seeded onto an ibidi flow chamber with 8 microelectrodes on the bottom (ECIS Flow Array 1E). Experiments proceeded exactly as ibidi flow experimental setup, with the addition of impedance readings taken every five min across each electrode for 72 hr.

### Statistical Analysis

All statistical analyses were performed using Prism v8.01 (www.graphpad.com), with an *α* of 0.05. For two-sample data sets with equal variances (control -v-a single experimental condition) unpaired, two-tailed Student’s *t*-test was used as reported in figure legends. For data sets with greater than two conditions and equal variances, one-way analysis of variance (ANOVA) with Tukey’s *post-hoc* test was used as reported in figure legends. * p ≤ 0.05, ** p ≤ 0.01, *** p≤ 0.001, **** p≤ 0.0001, ns, not significant.

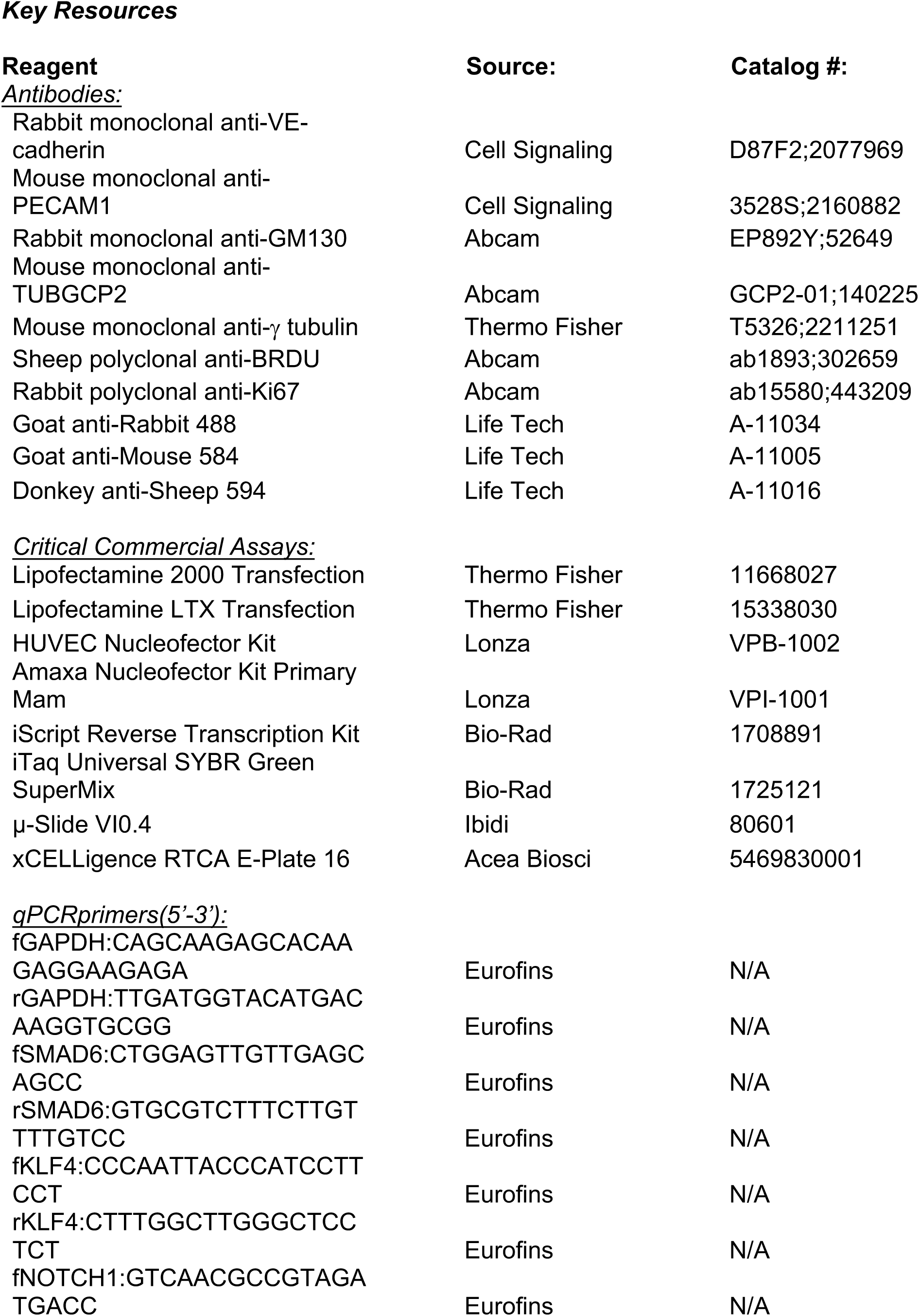
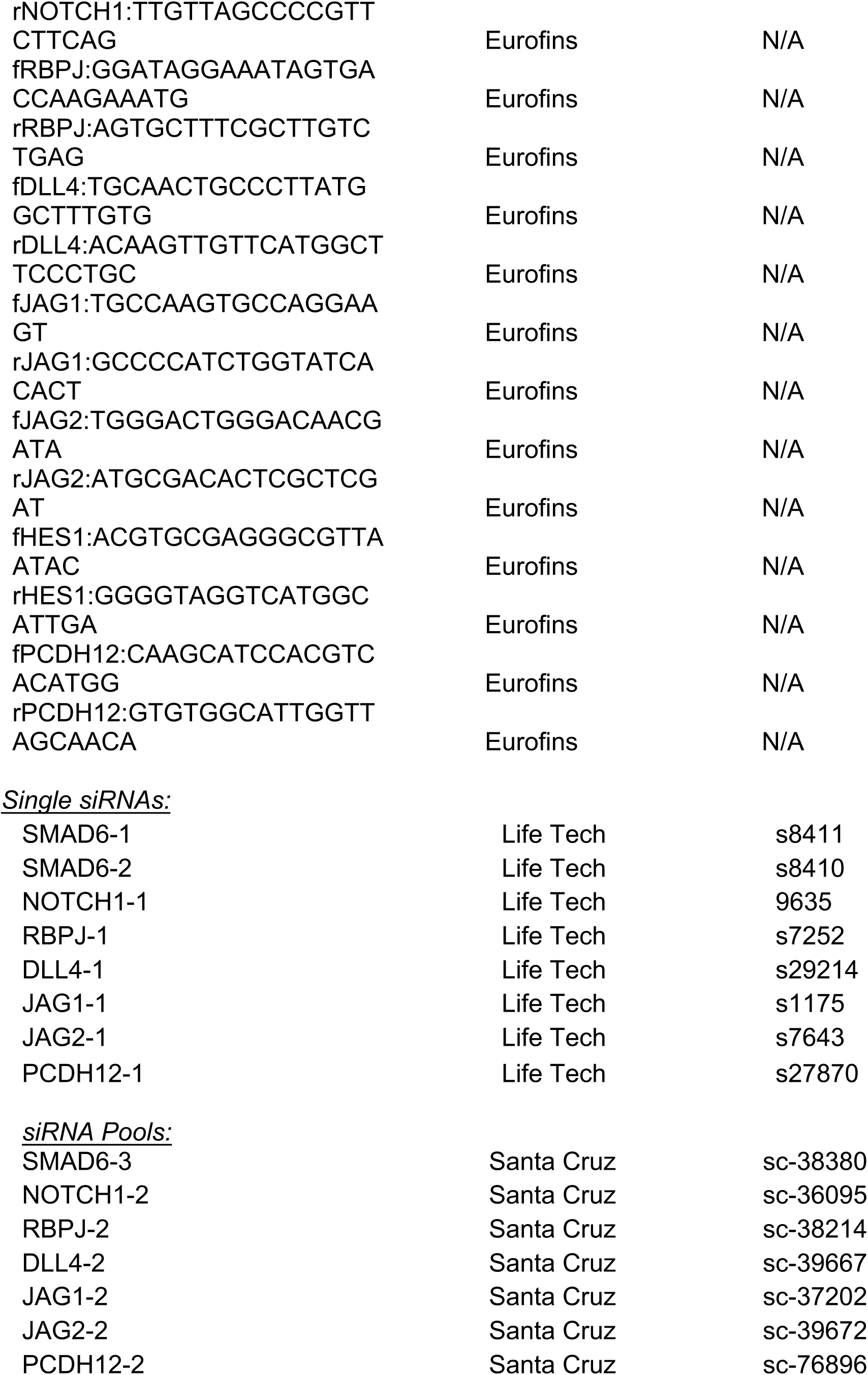

### Supplemental Files

The supplement includes 3 supplemental figures and figure legends.

## ACKNOWLEDGEMENTS

We thank Bautch lab lab members for input and constructive discussion, Dr. Keith Burridge for the use of TEER apparatus, and the UNC High Throughput Sequencing Facility (HTSF) for support. We thank Dr. Suk-Won Jin for constructive comments on the manuscript.

## AUTHOR CONTRIBUTIONS

Dana L Ruter (DLR) and Victoria L Bautch (VLB) conceptualized the work; DLR, Ziqing Liu (ZL), Kimlynn M Ngo (KMN), Shaka X (SX), and Elise J Kidder (EJK) performed and analyzed experiments; DLR and VLB wrote the manuscript; DLR, ZL, KMN, SX, Allison Marvin (AM), Danielle B Buglak (DBB) and VLB reviewed and edited the manuscript; VLB provided study supervision and oversight.

## SUPPLEMENTAL FIGURE LEGENDS

**Supplemental Figure 1(Linked to Fig 1).**
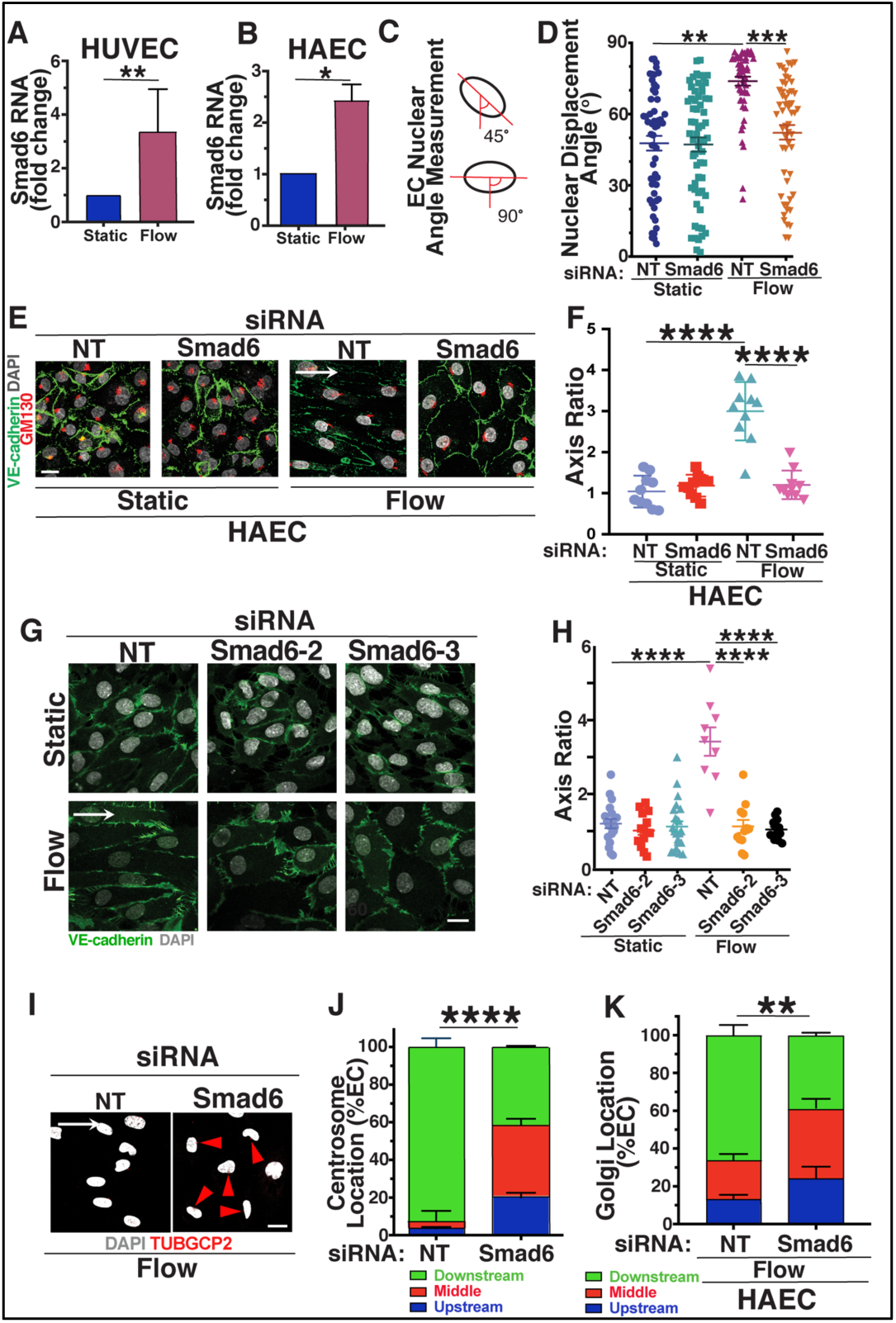
SMAD6 is Required for Homeostatic Endothelial Cell Flow-Mediated Alignment and Polarization. **A**,**B)** Smad6 qPCR RNA levels (normalized to static control) in HUVEC (A) and HAEC (B). Statistical analysis, Student’s t-test; *, p≤0.05; **, p≤0.01. **C)** Diagram of nuclear angle measurements used to quantify flow alignment. **D)** Nuclear angle quantification of **Fig1A. E)** Representative panels of HAEC stained with VE-cadherin (green, junctions), GM130 (red, Golgi), and DAPI (white, nucleus) under control (static) or flow conditions with indicated treatments. White arrow, flow vector. Scale bar, 20μM. **F)** Quantification of cell axis ratio of HAEC in indicated conditions from **Supp Fig 1E**. Statistical analysis, One-way ANOVA; ****, p≤0.0001. **G)** Representative panels of HUVEC with additional SMAD6 siRNAs (see Key Resources in Materials and Methods) stained with VE-cadherin (green, junctions) and DAPI (white, nucleus) under control (static) or flow conditions. White arrow, flow vector. Scale bar, 20μM. **H)** Quantification of cell axis ratio in indicated conditions from **Supp Fig 1G**. Statistical analysis, One-way ANOVA; ****, p≤0.0001. **I)** Representative panels of HUVEC stained with TUBGCP2 (red, centrosome) and DAPI (white, nucleus) under flow conditions with indicated treatments. White arrow, flow vector. Red arrowhead, centrosome location. Scale bar, 20μM. **J**) Quantification of centrosome location in **Supp Fig 1I** relative to nucleus in indicated conditions. n = ≥30 cells per condition. Statistical analysis, One-way ANOVA; ****, p≤0.0001. **K)** Quantification of HAEC Golgi localization relative to nucleus (**Supp Fig 1E**) in indicated conditions. n = ≥30 cells per condition. Statistical analysis, One-way ANOVA; **, p≤0.01.

**Supplemental Figure 2(Linked to Fig 1, Fig 2, Fig 3).**
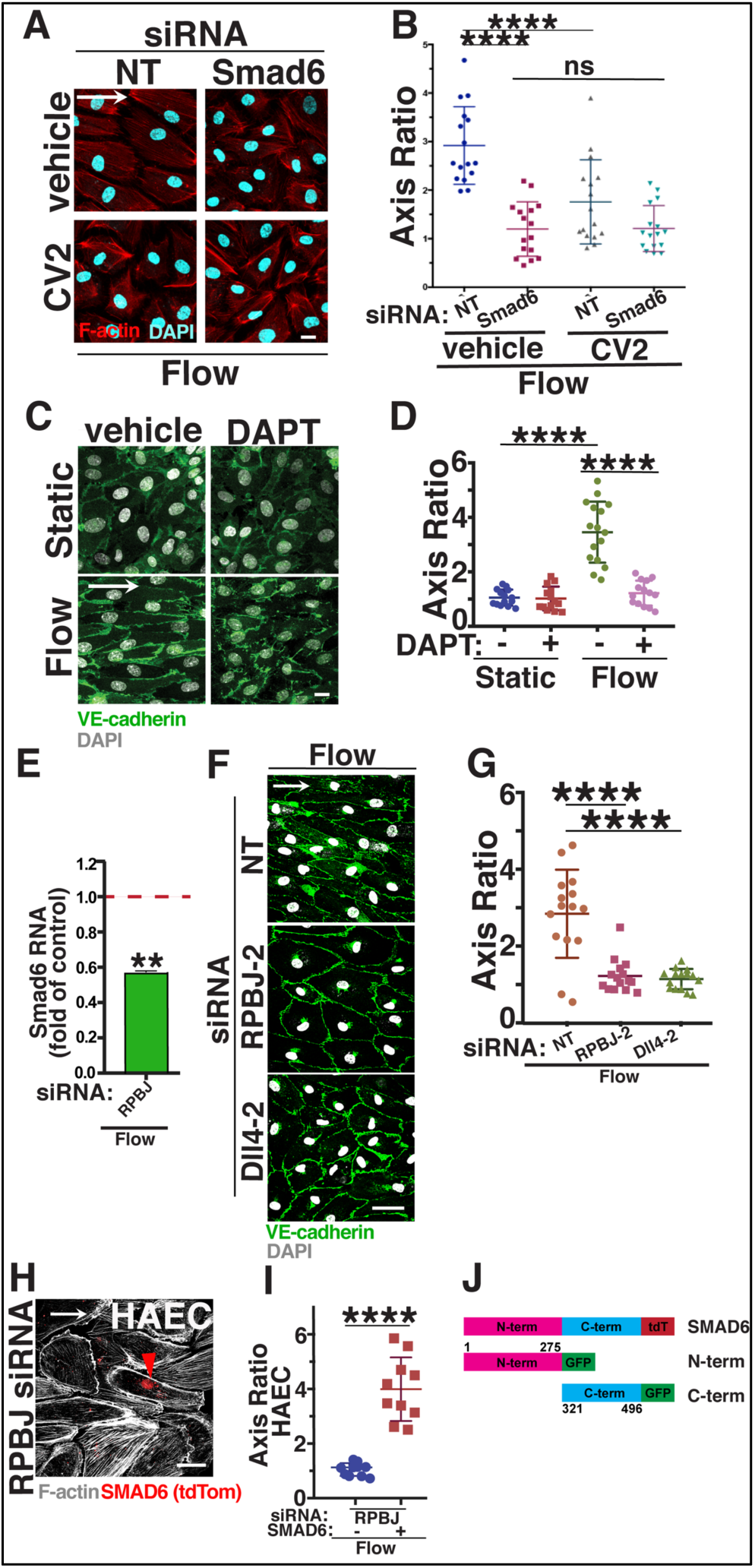
SMAD6 is Downstream of Notch Signaling in Homeostatic Endothelial Cell Flow-Mediated Alignment. **A)** Representative panels of HUVEC stained with Phalloidin (red, actin) and DAPI (white, nucleus) under flow conditions with indicated treatments. White arrow, flow vector. Scale bar, 20μM. **B)** Quantification of cell axis ratio in indicated conditions. Statistical analysis, One-way ANOVA; ****, p≤0.0001; ns, not significant. **C)** Representative panels of HUVEC stained with VE-cadherin (green, junctions) and DAPI (white, nucleus) under control (static) or flow conditions with indicated treatments. White arrow, flow vector. Scale bar, 20μM. **D)** Quantification of cell axis ratio in indicated conditions. Statistical analysis, One-way ANOVA; ****, p≤0.0001. **E)** qPCR RNA levels (normalized to static control) in HUVEC treated with RPBJ siRNA. Statistical analysis, Student’s t-test; **, p≤0.01 **F)** Representative panels of HUVEC with additional RBPJ and DLL4 siRNAs (see Key Resources) stained with VE-cadherin (green, junctions) and DAPI (white, nucleus) under control (static) or flow conditions. White arrow, flow vector. Scale bar, 50μM. **G)** Quantification of cell axis ratio in indicated conditions. Statistical analysis, One-way ANOVA; ****, p≤0.0001. **H)** Representative panel of HAEC treated with RPBJ siRNA and stained with phalloidin (F-actin, white) and expression construct (SMAD6) under flow conditions. White arrow, flow vector. Red arrowhead, positive EC. Scale bar, 20μM. **I)** Quantification of cell axis ratio in indicated conditions. Statistical analysis, student’s t-test; ****, p≤0.0001. **J)** Diagram showing SMAD6 constructs. Numbers indicate amino acids in human SMAD6.

**Supplemental Figure 3(Linked to Fig4).**
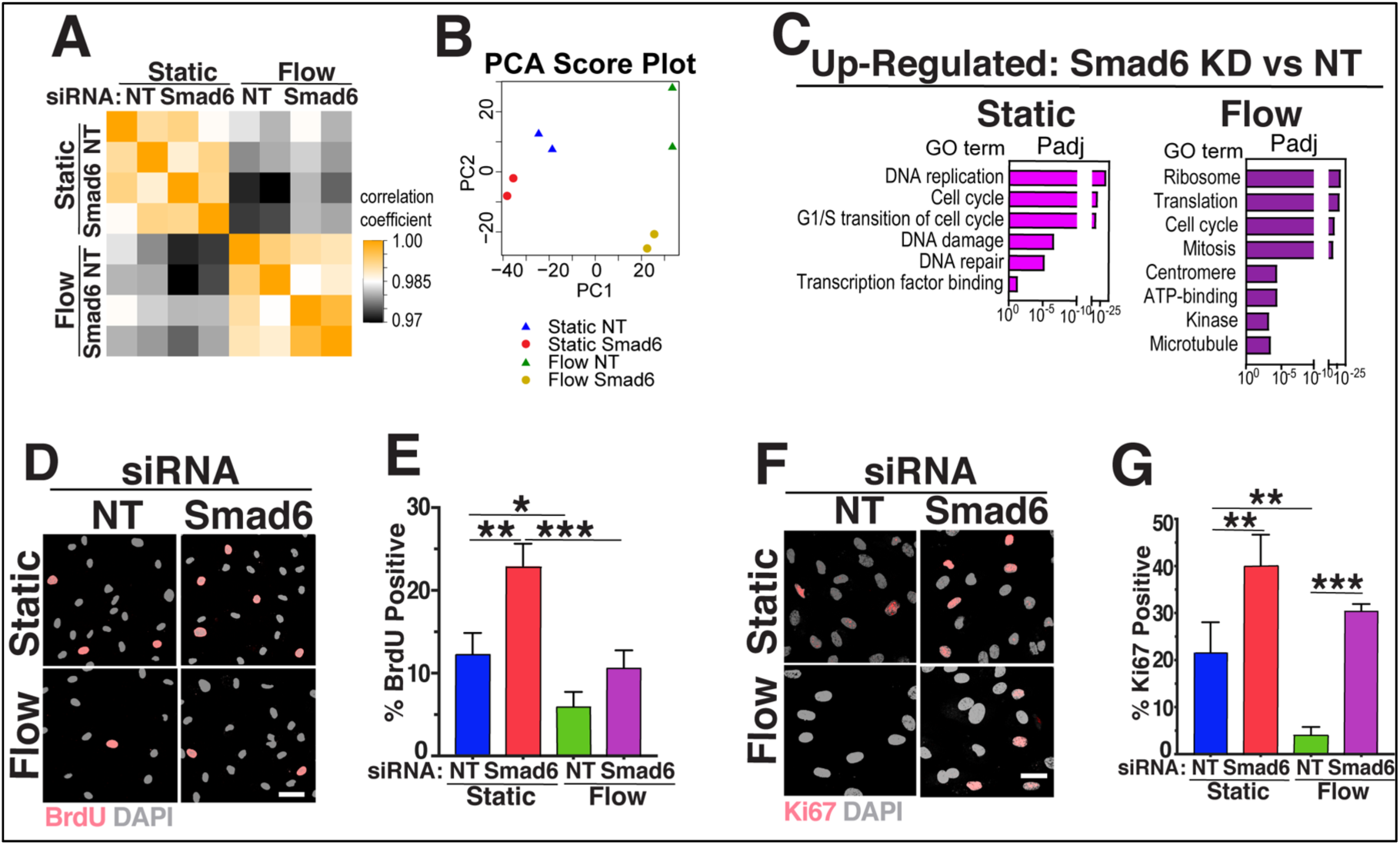
Endothelial Cell Transcriptomics Reveals Increased Proliferation with Reduced SMAD6 Levels. **A**) Pearson Correlation Analysis of bulk RNA-seq data from endothelial cells of indicated conditions. Correlation coefficients between any two samples plotted as heatmap.**B**) Principle Component (PC) Analysis of bulk RNA-seq data. **C**) GO analysis performed on differentially expressed genes from bulk RNA-seq data using DAVID. Representative GO terms significantly enriched (P adjusted < 0.1) in up-regulated genes are shown. **D**) Representative panels of HUVEC labeled with BrdU and stained with anti-BrdU (pink, S-phase marker) and DAPI (white, nucleus) under control (static) or flow conditions with indicated siRNA treatments. White arrow, flow vector. Scale bar, 20μM. **E)** Quantification of BrdU incorporation in indicated conditions. Statistical analysis, One-way ANOVA; *, p≤0.05; **, p≤0.01; ***, p≤0.001. **F)** Representative panels of HUVEC stained for Ki67 (pink, proliferation marker) and DAPI (white, nucleus) under control (static) or flow conditions with indicated siRNA treatments. White arrow, flow vector. Scale bar, 20uM. **G)** Quantification of Ki67+ cells in indicated conditions. Statistical analysis, One-way ANOVA; **, p≤0.01; ***, p≤0.001.

